# Lagrangian Formalism in Biology: I. Standard Lagrangians and their Role in Population Dynamics

**DOI:** 10.1101/2022.03.25.485848

**Authors:** D.T. Pham, Z.E. Musielak

## Abstract

The Lagrangian formalism is developed for the population dynamics of interacting species that are described by several well-known models. The formalism is based on standard Lagrangians, which represent differences between the physical kinetic and potential energy-like terms. A method to derive these Lagrangians is presented and applied to selected theoretical models of the population dynamics. The role of the derived Lagrangians and the energy-like terms in the population dynamics is investigated and discussed. It is suggested that the obtained standard Lagrangians can be used to identify physical similarities between different population models.

## 1. Introduction

The astounding progress of modern physics in discovering and understanding the fundamental laws of Nature that govern the structure and evolution of nonorganic matter has been caused by its powerful mathematical-empirical approach. In this approach, physical theories are formulated using the language of mathematics and their predictions are verified by experiments. At the same time, life sciences, included biology, remain mainly empirical and descriptive.

There have been attempts to formulate mathematical models of some biological systems, and thereby to establish mathematically oriented theoretical biology [1]. Different areas of mathematics have become increasingly more important in biology in recent decades, specifically, statistics in experimental design, pattern recognition in bioinformatics, and mathematical modeling in evolution, ecology and epidemiology [2]; however, it is pointed out that some of these attempts can be classified as ‘uses’ but others must be considered as ‘abuses’.

By combining theoretical and empirical biology, the mathematical-empirical approach may be established with expectations that its description of living organisms reaches the level of the description given by physics to nonorganic matter. In modern theoretical physics, all fundamental equations describing matter are derived by using the Lagrangian formalism [3-5], which requires a prior knowledge of functions called Lagrangians [6-8]. A number of different methods have been proposed [9-14] to obtain the Lagrangians for the most basic equations of modern physics [5].

In theoretical biology, Kerner [15] was first who applied the Lagrangian formalism to biology and obtained Lagrangians for several selected biological systems described by first-order ordinary differential equations (ODEs). Later, Paine [16] investigated the existence and construction of Lagrangians for similar set of ODEs following the original work of Helmholtz [6]. More recently, Nucci and Tamizhmani [17] derived Lagrangians for some models representing the population dynamics by using the method based on Jacobi Last Multiplier [10]. Moreover, Nucci and Sanchini [18] obtained Lagrangian for an Easter Island population model.

The main goal of this paper is to develop the Lagrangian formalism for biological systems. The formalism is based on standard Lagrangians, whose main characteristic is the presence of the difference between the kinetic and potential energy-like terms [5,9,10]. A method to derive these Lagrangians is presented and applied to selected models of the population dynamics. The role of the derived Lagrangians and the energy-like terms in these models is investigated and discussed. It is suggested that the obtained standard Lagrangians can be used to identify physical similarities between diverse biological systems.

The paper is organized as follows. Section 2 presents a brief overview of the Lagrangian formalism and standard Lagrangians. Section 3 describes and discusses the models of the population dynamics, and Section 4 concludes the paper.

## 2. Lagrangian formalism

### 2.1. General overview

Preliminary formulation of the Lagrangian formalism was originally done by Euler in 1742, and then it was refined and applied to Newtonian dynamics by Lagrange, who set up its currently used form in his *Analytic Mechanics* that first appeared in 1788; however, see [3] for more recent edition. According to Lagrange, the formalism deals with a functional 𝒮[*x*(*t*)], which depends on a continuous and differentiable function *x*(*t*) that describes evolution of a property of any dynamical system (given by *x*) in time (represented by *t*). This evolution is described by an equation of motion, which can be derived from the Lagrangian formalism.

The functional 𝒮 [*x*(*t*)] is called action and is defined by an integral over a scalar function *L* that depends on 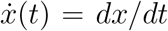, *x* and on *t*, so 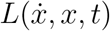 and it is called the Lagrangian function or simply Lagrangian. According to the principle of least action, or Hamilton’s principle [4,5], the functional 𝒮 [*x*(*t*)] must obey the following requirement *δ*𝒮 = 0, which says that the variation represented by *δ* must be zero to guarantee that the action is stationary (to have either a minimum, maximum or saddle point). The necessary condition that *δ*𝒮 = 0, is known as the Euler–Lagrange (E–L) equation that can be written as

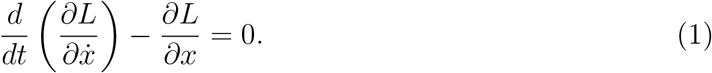

The Euler–Lagrange equation leads to a second-order ordinary differential equation (ODE) that can be further solved to obtain *x*(*t*) that makes the action stationary. The procedure forms the basis of calculus of variations, and it works well when the Lagrangian 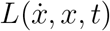 is already known. Deriving the second-order ODE from the E–L equation is called the *Lagrangian formalism*, and in this paper we deal exclusively with this formalism. It must be pointed out that the formalism has been extensively used in modern physics and that all its fundamental equations are derived by using it [5]. In this paper, we develop the Lagrangian formalism in biology and use it to obtain equations of motions for the selected models of population dynamics.

For dynamical systems whose total energy is conserved, the existence of Lagrangians is guaranteed by the Helmholtz conditions [6], which can also be used to obtain Lagrangians. The procedure of finding Lagrangians is called the inverse (or Helmholtz) problem of calculus of variations [7] and it shows that there are three separate classes of Lagrangians, namely, standard [3,4], nonstandard [8] and null [9] Lagrangians. Both standard and nonstandard Lagrangians give the same equations of motion after they are substituted into the E-L equation. However, null Lagrangians satisfy the E-L equation identically and therefore they do not give any equation of motion.

Our main goal is to establish the Lagrangian formalism for known ODEs that describe time evolution of different models of the population dynamics and find their Lagrangians; in this paper, we concentrate exclusively on standard Lagrangians.

### 2.2. Standrard Lagrangians

According to Lagrange [3], standard Lagrangians (SLs) are represented by differences between kinetic and potential energies of dynamical systems. In dynamics, if an oscillatory system of mass *m* is displaced by *x* in time *t*, then its kinetic energy of oscillations is given by

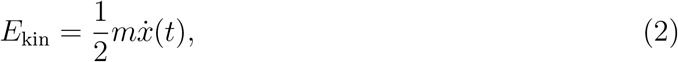

where 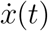 is the time derivative of the dynamical variable *x*(*t*), or otherwise known as 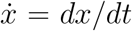. The potential energy depends on a spring constant *k* of the oscillatory system and it can be written as

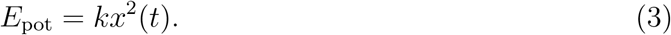

Thus, the standard Lagrangian for this system is

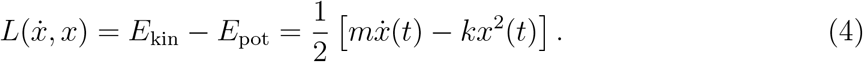

Substitution of this Lagrangian into the E-L equation (see Eq. 1) gives the following equation of motion:

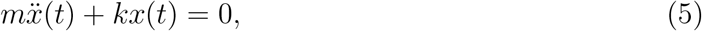

where 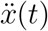 is the second derivative of *x*(*t*) with respect to time, or 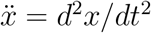.

The presented above form of the SL is characteristic for oscillatory systems that are not damped and not driven [2]. Obviously, the meaning of the variable *x*(*t*) may change from one physical system to another. Moreover, the meaning of *x*(*t*) in biological systems will be different than in physical systems, so will the meaning of the constants *m* and *k*. Nevertheless, a Lagrangian with a kinetic-like energy term, which contains 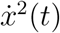, and a potential-like energy term, which contains *x*^2^(*t*), will be identified as the SL. The main objective of this paper is to derive SLs for different models of the population dynamics and discuss the role of these SLs in mathematical description of these models.

The Lagrangian formalism based on standard Lagrangians have been well-established in most fields of modern physics (e.g., [5] and references therein). Specifically, the formalism is commonly used to obtain equations of motion for dynamical systems in Classical Mechanics [3,7,8]. More recently, several methods were developed to solve the inverse Helmholtz problems for physical systems described by ODEs (e.g., [10-14]). Some of these methods developed for physical systems will be used in this paper to derive Lagrangians for biological systems.

As already mentioned in the Introduction, there have been several attempts to establish the Lagrangian formalism in biology [15-18], and different Lagrangians were obtained for some selected biological systems. In this paper, we concentrate on standard Lagrangians (SLs) because of their specific physical meaning discussed above. Our goals are to derive SLs for some of the model for population dynamics, and discuss their meaning and role of these Lagrangians in biology.

### 2.3. Method to derive standard Lagrangians

The Lagrangian formalism requires prior knowledge of a Lagrangian. In general, there are no first principle methods to obtain Lagrangians, which are typically presented without explaining their origin. In physics, most dynamical equations were established first and only then their Lagrangians were found, often by guessing. Once the Lagrangians are known, the process of finding the resulting dynamical equations is straightforward and it requires substitution of these Lagrangians into the E-L equation. There has been some progress in deriving Lagrangians for physical systems described by ODEs (e.g., [10-14]), and in this paper, we develop a new method to obtain standard Lagrangians (SLs) for several models that describe interacting species of the population dynamics. Let us point out that the same method can be used in physics and in other natural sciences to obtain Lagrangians for systems described by second-order ODEs.

The main objective is to solve the inverse (Helmholtz) problem of the calculus of variations [6,7] and derive the standard Lagrangian for a given second-order ODE. Let us consider the following ODE

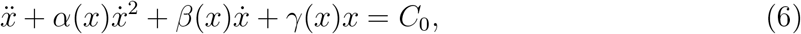

where *α*(*x*), *β*(*x*) and *γ*(*x*) are at least twice differentiable functions of the dependent variable only, and these functions are to be specified by the selected models of the population dynamics. In addition, *C*_0_ is a constant that may appear in some of these models (see Section 3). From a physical point of view, the above equation of motion describes an oscillatory system that is affected by two damping terms with *α*(*x*) and *β*(*x*), and driven by the constant force *C*_0_. In case, *α*(*x*) = *β*(*x*) = *C*_0_ = 0, the equation represents a harmonic oscillator [4,5].

Let us consider a simplified form of this ODE by taking *β*(*x*) = 0 and *C*_0_ = 0, and obtain

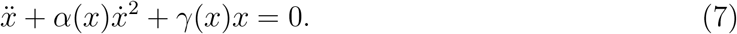

Despite the presence of the damping-like term 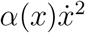, the equation is considered to be conservative, which means that its standard Lagrangian can be obtained [11,20] and written as

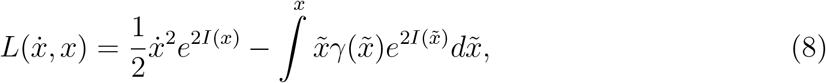

with

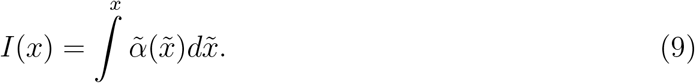

The main reason for reducing Eq. (6) to Eq. (7) is that the term 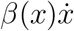 is by itself a null Lagrangian that identically satisfies the E-L equation [21-23], therefore, its derivation from any SL is not possible [11,12]. In the following, we propose to account for this term in a novel way.

We write Eq. (7) in the following form

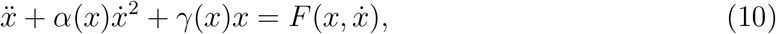

where the force-like term becomes

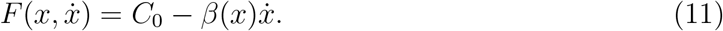

All population dynamics models considered in this paper (see Section 3) are represented by second-order ODEs in the form of Eq. (10) with 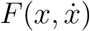 given by Eq. (11).

Our results presented in the Appendix generalize the previous work [20] and demonstrate that the Lagrangian given by Eq. (8) is also the SL for Eq. (10) if, and only if, this SL is substituted into the E-L equation

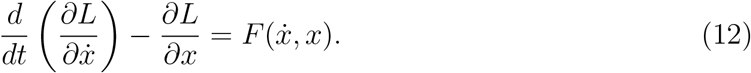

This form of the E-L equation is commonly known in physics, and the presence of the force-like term on the right-hand side of the equation is justified by 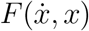 being a force that does not arise from a potential (e.g., [4]); for further discussion of this term, see Section 3.3.

It is straightforward to verify that substitution of Eq. (8) into Eq. (12) gives the original equation given by Eq. (10). This makes our method easy to find SLs for any ODEs of the form of Eq. (10) as it requires identifying the functions *α*(*x*), *γ*(*x*) and 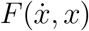, and evaluating the integral in Eq. (9) as well as the one in the SL given by Eq. (8).

Our method is simple and easy to use for the population dynamics models considered in this paper. Let us point out that this method can be applied to any dynamical system that is described by the second-order ODEs of the form of Eq. (10) in any areas of natural sciences.

## 3. Applications to models of population dynamics

### 3.1. Selected models

The models of the population dynamics considered in this paper are listed in Table 1. Our selection process was guided by the previous work, by Trubatch and Franco[17], and Nucci and Tamizhmani [18]. Both papers considered the well-known population models that involve two interacting species described by coupled nonlinear ODEs, namely, the Lotka-Volterra, Gompertz, Verhulst and Host-Parasite models as shown in Table 1. The authors of these papers determined Lagrangians corresponding to the ODEs representing mathematically the models, either *ad hoc* [19] or using the method of Jacobi Last Multiplier [17]; the Lagrangians of the same form were obtained, and they were treated as the generating functions for the ODEs representing the models [17,18].

| Population models | Equations of Motion |
| --- | --- |
| Lotka-Volterra Model | $\dot{w}_1 = w_1(a + bw_2)$<br>$\dot{w}_2 = w_2(A + Bw_1)$ |
| Verhulst Model | $\dot{w}_1 = w_1(A + Bw_1 + f_1w_2)$<br>$\dot{w}_2 = w_2(a + bw_2 + f_2w_1)$ |
| Gompertz Model | $\dot{w}_1 = w_1(A \log\left(\frac{w_1}{m_1}\right) + Bw_2)$<br>$\dot{w}_2 = w_2(a \log\left(\frac{w_2}{m_2}\right) + bw_1)$ |
| Host-Parasite Model | $\dot{w}_1 = w_1(a - bw_2)$<br>$\dot{w}_2 = w_2(A - B\frac{w_2}{w_1})$ |
| SIR Model | $\dot{w}_1 = -bw_1w_2$<br>$\dot{w}_2 = bw_1w_2 - aw_2$ |

As it is well-known, Lagrangians can be of different forms and yet they would give the same equation of motion [5,7]. In most cases, the forms of these Lagrangians do not resemble the SLs in which the kinetic and potential energy-like terms can be identified [10-14]. However, the main objective of this paper is to derive the SLs and for some selected models of the population dynamics and compare the obtained SLs to those previously found [18,19]. Because of the specific physical meaning of the SLs derived here, we are able to address the role and meaning of these SLs in the population dynamics.

In selecting models of the population dynamics, we used the four models used in the previous studies [19,18]. In addition, we selected the SIR model (see Table 1).

The first four models of the population dynamics presented in Table 1 describe two interacting species (preys and predators) of the respective populations *w*_1_(*t*) and *w*_2_(*t*) that evolve in time *t*, which is denoted by the time derivatives 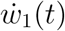 and 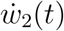. The coefficients *a, A, b, B f*_1_, *f*_2_, *m*_1_ and *m*_2_ are real and constant parameters that describe the interaction of the two species. The Lotka-Volterra, Verhulst and Gompertz models are *symmetric*, which means that the depedent variables can be replaced if, and only if, the constants are replaced, *a* → *A, b* → *B, f*_1_ → *f*_2_ and *m*_1_ → *m*_2_. However, the Host-Parasite model is *asymmetric* in the dependent variables.

The SIR model presented in Table 1 describes the spread of a disease in a population and the dependent variables *w*_1_(*t*) and *w*_2_(*t*) represent susceptible and infectious populations, with *a* and *b* being the recovery and infection rates, respectively. Similarly to the Host-Parasite model, the SIR model is also *asymmetric* but the origin and nature of this asymmetry in both models is significantly different.

### 3.2. Steps to derive standard Lagrangians

We solve the inverse (Helmholtz) calculus variational problem and derive standard Lagrangians for all selected models of the population dynamics shown in Table 1. The following steps must be undertaken to solve the problem:

1. Convert the set of coupled nonlinear ODEs for each model into a second-order nonlinear ODE for each dependent variable.
2. Cast the derived second-order ODEs into the equation of the same form as Eq. (10).
3. Compare the obtained second-order ODEs to Eq. (10) and identify the functions *α*(*x*), *γ*(*x*) and 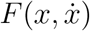 for each variable in each model.
4. Evaluate the integral in Eq. (9) and then determine the exponential factor.
5. Evaluate the integral in Eq. (8).
6. Use the above integrals and Eq. (8) to find the standard Lagrangian for each variable in each model.
7. Verify the derived standard Lagrangian by substituting it into the Euler-Lagrange equation given by Eq. (12).

As pointed out in [19], the choice between the first and second order description of the population dynamics is typically motivated by the information available. In case only one population is observed, then the second-order ODE can be solved for this population. To make it more general, we derive the second-order ODE for each variable in each model, so both equations can be solved if both populations are observed. For this paper and the presented results, the second-order ODEs are required because our method to derive the SLs is valid only for such equations.

Our method to derive the SLs for the selected models is straightforward and the above steps are easy to perform and they always give the standard Lagrangian for any second-order ODEs of the same form as Eq. (10). In the following, we present the derived second-order ODEs for each variable of the model and the resulting SLs.

### 3.3. Standard Lagrangians for selected models

Our method to derive standard Lagrangians for the models presented in Table 1 requires that the systems of coupled nonlinear first-order ODEs are cast into one second-order ODE for each variable. All derived second-order ODEs can be expressed in the same form as Eq. (10), which can be written as

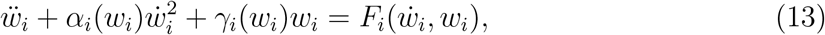

where *i* = 1 and 2. Since *w*_*i*_(*t*) represents the population of species, its derivative with respect of time 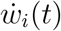 describes the rate with which the population changes, and 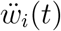 its acceleration. Despite the presence of the damping-like term 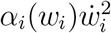, the LHS of the above equation is conservative [11,20] and it describes oscillations of the population of species with respect to its equilibrium. These oscillations are modified by the force-like term on the RHS of the equation. Let us now describe this term.

Typically, the presence of any term with 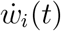 corresponds to friction forces in classical mechanics. In the approach presented in this paper, all friction-like terms that explicitly depend on 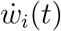 are collected on the RHS of the equation as 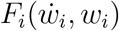, which becomes the force-like term. Since 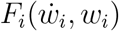 arises directly from the friction-like terms, its origin is not potential, and therefore this force-like term may appear on the RHS of the E-L equation (see Eq. 12) as it is shown in [4].

In our derivations of the standard Lagrangians for the models of the population dynamics presented in Table 1, we follow the above steps. As an example, we show the calculations required for each step for the Lotka-Volterra model. However, for the remaining four models in Table 1, we only present the final results.

## 4. Lotka-Volterra Model

The Lotka-Volterra model was developed by chemist Alfred Lotka in 1910 and mathematician Vito Volterra in 1926 [24,25], and this model describes the interaction of two populations (predator-prey) based on the assumptions that the prey increases exponentially in time without the predator, and the predator decreases exponentially without the prey [19]. The model is symmetric and it is represented mathematically by a system of coupled nonlinear ODEs given in Table 1.

*Step 1:* Using the coupled nonlinear ODEs for this model, we convert the equations into the following second-order ODEs for the variables *w*_1_ and *w*_2_

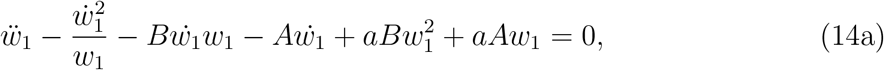

and

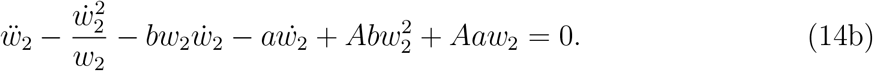
*Step 2:* We cast these equations into the form of Eq. (13), and obtain

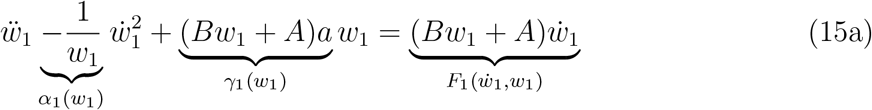

and

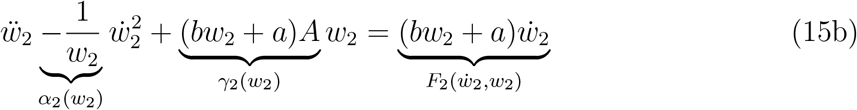
*Step 3:* Using the above equations, we identify the functions

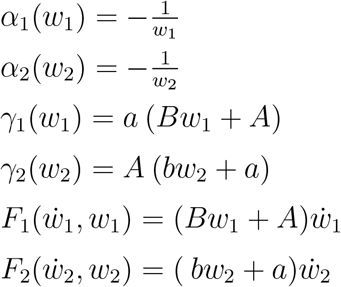
*Step 4:* Having identified *α*(*w*_1_) and *α*(*w*_2_), the intergral given by Eq. (9) can be evaluated, and the result is

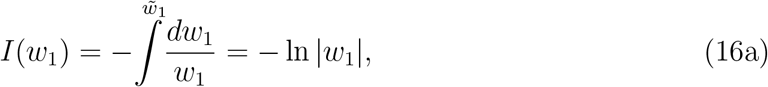

and

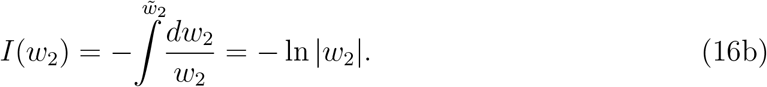 Then, the factors 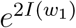 and 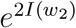 become

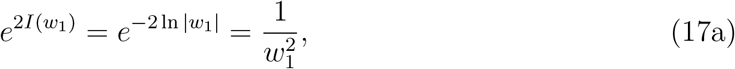

and

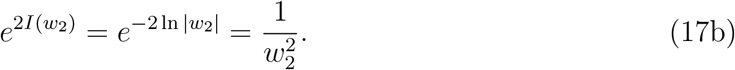
*Step 5:* Since *γ*(*w*_1_) and *γ*(*w*_2_) are known, the intergral in Eq. (8) can be calculated, and we find

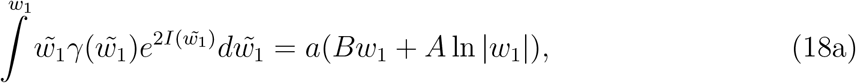

and

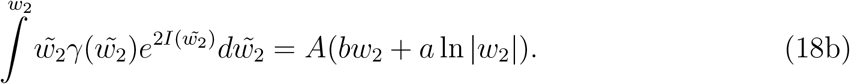
*Step 6:* Using the above results and the definition of Lagrangian given by Eq. (8), the following standard Lagrangians for the two dependent variables of the Lotka-Volterra model are obtained:

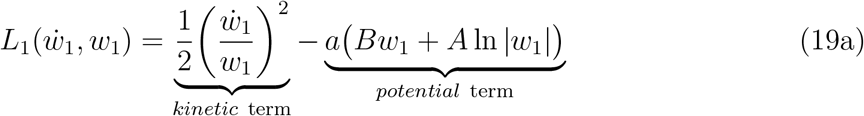

and

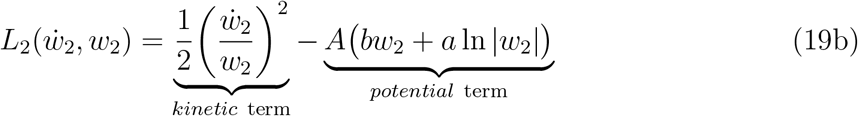
*Step 7:* Substituting the derived standard Lagrangians and 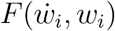 into the following Euler-Lagrange equations

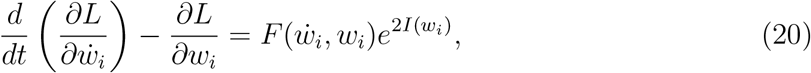

where *i* = 1 and 2, we obtain Eqs. (15a) and (15b). This verifies that the presented method to derive the SLs is valid.

## 5. Verhulst Model

This logistical (or Verhulst-Pearl) equation was first introduced by the Belgian statistician Pierre Francois Verhulst in 1838 [27]. This model describes the organisms’ growth dynamics in a habitat of finite resources, which means the population is limited by a carrying capacity. This model is valueable for optimisation of culture media by developing strategies and selection of cell lines.

In this paper, the Verhulst model describes the population of interacting species by considering self-interacting terms that prevent the exponential increase or decrease in the size in the populations observed in the Lotka-Volterra model [19]. The system of coupled nonlinear ODEs given in Table 1 shows that this model is symmetric.

After performing steps 1 through 3, the following functions for this model are obtained:

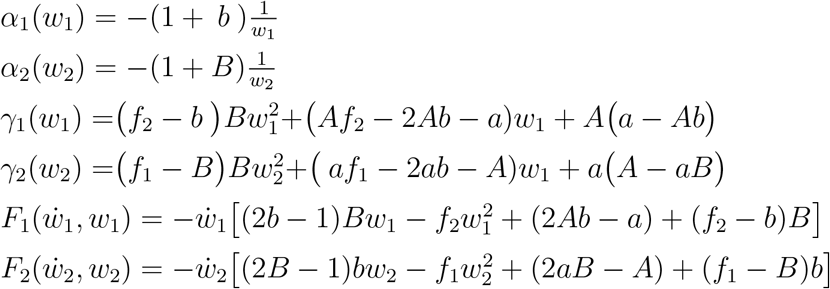

Substitution of these functions into Eq. (13) gives the second-order ODEs for the variables *w*_1_ and *w*_2_.

Then, we implement steps 4 through 6 and the resulting standard Lagrangians for the variables *w*_1_ and *w*_2_ of the Verhulst model are

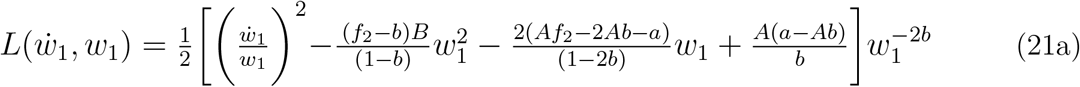

and

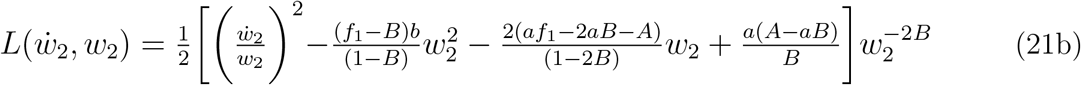

The kinetic and potential energy-like terms are easy to recognize, and the functions 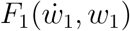 and 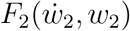 are given above. Substitution of these Lagrangians into the E-L equations (see Eq. 20) validates the method.

## 6. Gompertz Model

English economist Benjamin Gompertz proposed a model to describe the relationship between increasing death rate and age in 1825. This model is useful in describing the rapid growth of a certain population of organisms such as the growth of tumors [28]. As well as, modelling the amount of medicine in the bloodstream.

Here, we follow [17,19] and consider the Gompertz model for the population dynamics This model generalizes the Lotka-Volterra model by including self-interaction terms that prevent an unbounded increase of any isolated population [19]; the self-interacting terms in the Gompertz model are different than those in the Verhulst model. The mathematical representation of this model given by the coupled and nonlinear ODEs in Table 1 shows that the model is symmetric.

The steps 1 through 3 allow us to identify the following functions in Eq. (13)

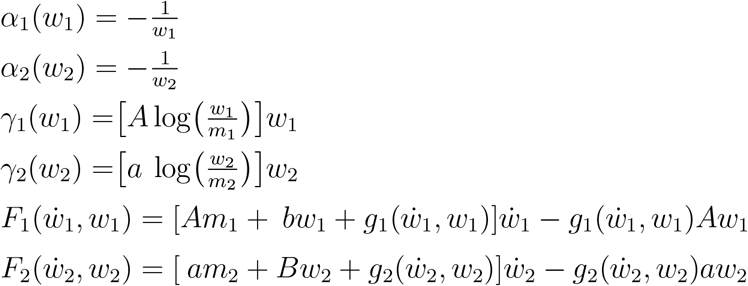

where

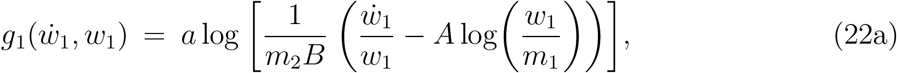

and

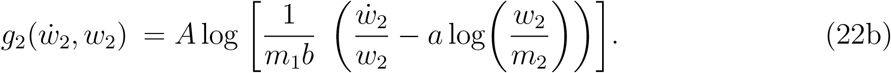

Then, the remaining steps result in the following standard Lagrangians for the two dependent variables of the Gompertz model

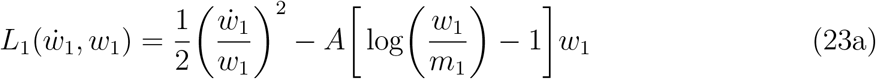

and

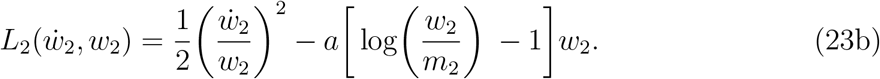

In both Lagrangians the kinetic and potential energy-like terms are seen, and the forcing functions 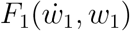 and 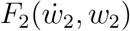 are given above. If we substitute these Lagrangians into Eq. (20), the second-order ODEs for the variables *w*_1_ and *w*_2_ are obtained.

## 7. Host-Parasite Model

This model describes the interaction between a host and its parasite. The model takes into account nonlinear effects of the host population size on the growth rate of the parasite population [19]. The system of coupled nonlinear ODEs (see Table 1) is asymmetric in the dependent variables *w*_1_ and *w*_2_.

Using steps 1 through 3, we find

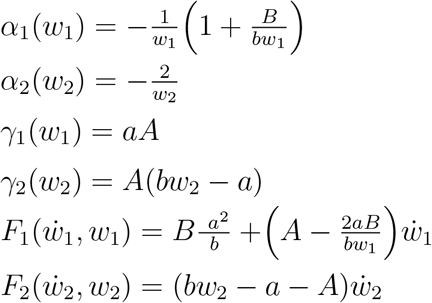

After substituting these functions into Eq. (13), the second-order ODEs for the variables *w*_1_ and *w*_2_ are obtained, and the equations are asymmetric, which means that the remaining steps 4 through 6 must be applied to each dependent variable (*w*_1_ or *w*_2_) separately.

The standard Lagrangian for the variable *w*_1_ is

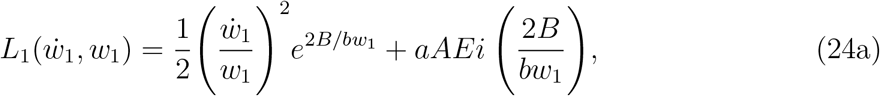

where the exponential integral *Ei*(2*B/bw*_1_) is a special function defined as

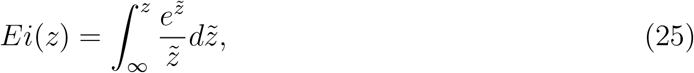

with *z* = 2*B/bw*_1_. It must be noted that *Ei*(*z*) is not an elementary function. Now, the standard Lagrangian for the variable *w*_2_ is given by

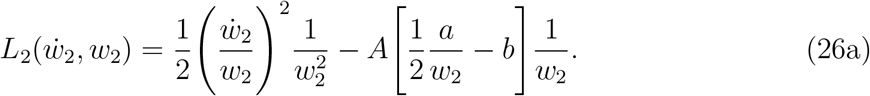

It is seen that there are significant differences between the Lagrangian for *w*_2_ and that for *w*_1_ in both the kinetic and potential energy-like terms. The differences are especially prominent in the potential energy-like terms, whose explicit dependence on the exponential integral *Ei*(2*B/bw*_1_) is a new phenomenon. The differences are caused by the asymmetry between the dependent variables in the original equations (see Table 1), which makes this model different than the fully symmetric Lotka-Volterra, Verhulst and Gompertz models, whose standard Lagrangians are also fully symmetric.

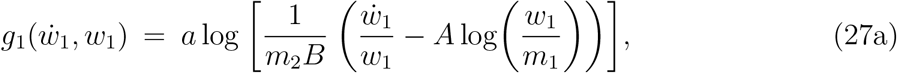

and

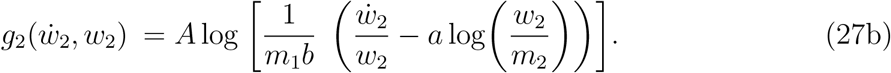

## 8. SIR Model

Kermack and McKendrick in 1927 derived the system of the first ODEs (see Table 1) describing the spread of a disease in a population [29]. It is the one of the simplest model, dividing the population into three distinct sub-populations: a susceptible population denoted by *w*_1_(*t*), the infectious population represented by *w*_2_(*t*), and a recovered population, we denote as *w*_3_(*t*).

It is seen that the dependent variable *w*_3_(*t*) does not appear explicitly in the set of ODEs given in Table 1 because it is related to *w*_1_(*t*) and *w*_2_(*t*) through the following population conservation law: *d/dt*(*w*_1_ +*w*_2_ +*w*_3_) = 0, which means that the sum of the three populations must remain constant in time. Moreover, *a >* 0 is the recovery rate and *b >* 0 is the rate of infection, which means that the terms *−bw*_1_*w*_2_ and *−aw*_2_ represent newly infected and recovered individuals, respectively.

After performing steps 1 through 3, we obtain

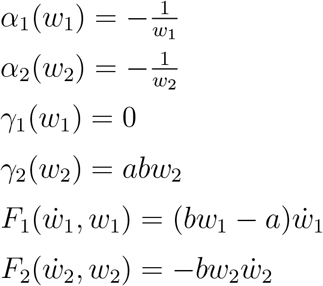

Then, steps 4 through 6 give the following standard Lagrangians

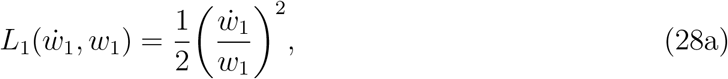

and

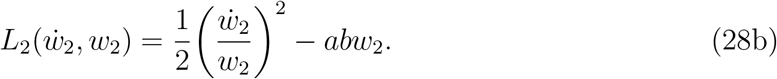

The fact that the SIR model is asymmetric is shown the lack of the potential energy-like term in 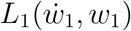 and its presence in 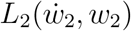. However, the kinetic energy-like terms are the same for the SLs for both variables, and they are also similar to such terms in the SLs obtained for the other population dynamics models.

### 8.1. Comparisons of Lagrangians and models

The derived standard Lagrangians for the considered models of the population dynamics are characterized by the terms corresponding to the kinetic and potential energy as well as to the forcing function. These terms are easy to identify (see Eqs. 19a and 19b) and they can be used to make comparisons between the Lagrangians and models they represent. The models considered in this paper can be divided into two families, namely, symmetric (Lotka-Volterra, Verhulst and Gompertz) and asymmetric (Host-Parasite and SIR) models. The SLs derived for these models are different than the Lagrangians previously obtained [19,17]; the main difference is the explicit time-dependence of those Lagrangians as compared to the SLs derived in this paper. In the following, we describe the general characteristics of the derived SLs and comment on their explicit time-independence.

The kinetic energy-like terms in all four models have the same factor 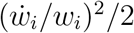, where *I* = 1 and 2, which represents the ratio at which the population changes to its value at a given time. For the Verhulst and Host-Parasite models, this ratio is modified by the other factors that depend on the concentration of species at a given time. It is interesting that the kine tic energy-like terms in the Lotka-Volterra, Gompertz and SIR models are independent from any constant parameters but for the other two models they are; in case of the Host-Parasite models only the variable *w*_1_ shows such a dependence.

The potential energy-like terms of the Lotka-Volterra model depends linearly on the concentration of species; however, the Verhulst, Gompertz and Host-Parasite models also have nonlinear (second-order) terms in the concentration of species. The SIR model is exceptional as its SL for the variable *w*_1_ does not depend on any potential energy-like term. On the other hand, the SL for the variable *w*_2_ does depend on the potential energy-like term that is linear in this variable. In all models, the potential energy-like terms depend on the constant parameters that appear in the derived second-order ODEs for these models. An interesting result is the presence of logarithmic terms in the Lotka-Volterra and Gompertz models, and the exponential integral *Ei* for the variable *w*_1_ for the Host-Parasite model. It must be also noted that the form of the potential energy-like term for the SIR model is the simplest among all the models considered here.

Since the kinetic and potential energy-like terms depend on the square of the rate with which the populations change and the square of their concentration, respectively, one may conclude that the derived SLs describe oscillatory systems. The fact that none of the obtained SLs depends explicitly on time indicates that the considered models oscillate in time with certain frequencies around the equilibrium, and that these oscillations are not directly affected by any ‘physical damping’ associated with the presence of terms that depend on 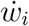, which are absent in derived the SLs; this is the main difference between the SLs of this paper and those previously obtained [19,17].

However, we must keep in mind that all damping terms represented by 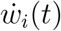 have been moved to the force-like functions denoted by 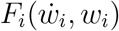, which significantly vary for different models. First, let us point out that the force-like functions become null Lagrangians [21-23], which means that they identically satisfy the E-L equation, and therefore they cannot contribute to any equation of motion. As a result, no standard Lagrangian can properly account for them [11,12] because the presented Lagrangian formalism is valid only for conservative systems.

Second, the force-like functions may depend only on 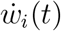 and on 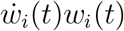, and the constant parameters, or may depend on higher powers of thess variables that are arguments of the logarithmic functions. As the presented results demonstrate, the forms of the force-like functions significantly differ for different models, with the simplest being for the SIR and Lotka-Volterra models, and then the increasing complexity for the Host-Parasite and Verhulst models. The most complex form of 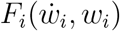 is found for the Gompertz model.

Third, the observed increase in complexity of the force-like function is caused by the role played by the terms that depend on 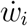 as well as on the combination of terms with 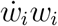. In other words, the forms of 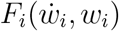 significantly affect the oscillatory behavior of the systems represented by the SLs (see discussion above). The force-like function may modify this behavior by causing the systems to reach the equilibrium faster or diverge from it; detailed analysis requires solutions to the ODEs representing the considered systems, which is out of the scope of this paper.

Finally, let us point out that since the energy-like terms in the SLs have specific meanings for the models, we suggest that they may be used to identify physical similarities between different models as well as they may be utilized to classify the models into categories that have similar biological characteristics. Despite the fact that the SLs were derived only for the population dynamics models, the developed method and the above discussion can be easily applied to a broad range of biological systems, which will be explored in succeeding papers.

## 9. Conclusions

We developed Lagrangian formalism for the following population dynamics models: Lotka-Volterra, Verhulst, Gompertz, Host-Parasite and SIR models. For ODEs that represent these models, we solved the inverse (Helmholtz) variational calculus problem and derived standard Lagrangians for the models. The main characteristic of these Lagrangians is that their kinetic and potential energy-like terms are identified and that they can be used to make comparisons between the obtained Lagrangians as well as the models.

The comparison demonstrates the role of these terms in the population dynamics models and gives new insights into the models by showing their similarities and differences. Moreover, the analogy between the derived standard Lagrangians and that known for a harmonic oscillator in physics is used to discuss the oscillatory behavior of the models with respect to their equilibrium. In our approach, we collected the terms with the first-order derivatives in time and identified them as the force-like functions, which happened to be significantly different for each model and strongly depend on the parameters of each model. By separating the force-like functions, we were able to see the effects of these functions on the model’s oscillatory behavior.

Our method of solving the inverse calculus of variation problem and deriving standard Lagrangians is applied to the models of population dynamics. However, the presented results show that the method can be easily extended to other biological systems whose equations of motion are known.

## References

[1] D. Krakauer, J.P. Collins, D. Erwin, J.C. Flack et al., ”The challenges and scope of theoretical biology”, J. Theor. Biology, 276, 269–276, 2011.

[2] R.M. May, ”Uses and abuses of mathemcatics in biology”, Science, 303, 790–793, 2011.

[3] Lagrangian, J.L. Analytical Mechanics, Springer, Dordrecht, The Niederlands, 1997.

[4] H. Goldstein, C. Poole, J. Safko, Classical Mechanics, Addison-Wesley, New York, 2002.

[5] J.V. José, E.J. Saletan, Classical Dynamics, A Contemporary Approach, Cambridge Univ. Press, Cambridge, 2002.

[6] H. Helmholtz, J. f. d. Reine u. Angew Math., 100 (1887) 213

[7] J. Lopuszanski, The Inverse Variational Problems in Mechanics, World Scientific, 1999.

[8] V.I. Arnold, Mathematical Methods of Classical Mechanics, Springer, New York, 1978

[9] P.J. Olver, Applications of Lie Groups to Differential Equations, Springer-Verlag, New York, 1993.

[10] M.C. Nucci and P.G.L. Leach, ”Lagrangians galore”, J. Math. Phys., 48, 123510, 2007.

[11] Z.E. Musielak, ”Standard and non-standard Lagrangians for dissipative dynamical systems with variable coefficients”, J. Phys. A Math. Theor., 41, 055205, 2008.

[12] J.L. Cieśliński and T. Nikiciuk, ”A direct approach to the construction of standard and non-standard Lagrangians for dissipative-like dynamical systems with variable coefficients”, J. Phys. A Math. Theor., 43, 175205, 2010

[13] R.A. El-Nabulsi, ”Fractional action cosmology with variable order parameter”, Int. J. Theor. Phys., 56, 1159, 2017.

[14] Z.E. Musielak, N. Davachi and M. Rosario-Franco, ”Special functions of mathematical physics: A unified Lagrangian formalism”, Mathematics, 8, 379, 2020.

[15] E.H. Kerner, ”Dynamical aspects of kinetics”, Bulletin of Mathematical Biophysics, 26, 333–349, 1964.

[16] G.H. Paine, ”The development of Lagrangians for biological models”, Bulletin of Mathematical Biology, 44, 749–760, 1982.

[17] M.C. Nucci and K.M. Tamizhmani, ”Lagrangians for Biological Models: The Method of Jacobi Last Multiplier”, J. Nonlinear Math. Phys., 19, 12500021, 2012.

[18] M.C. Nucci and G. Sanchini, ”Symmetries, Lagrangians and conservation laws of an Easter Island population model”, Symmetry, 7, 1613–1632, 2015.

[19] S.L. Trubatch and A. Franco, ”Canonical Procedures for Population Dynamics”, J. Theor. Biol, 48, 299–324, 1974.

[20] Z.E. Musielak, D. Roy and L.D. Swift, ”Method to derive Lagrangian and Hamiltonian for a nonlinear dynamical system with variable coefficients”, Chaos, Solitons Fractals, 38, 894, 2008.

[21] P.J. Olver, Applications of Lie Groups to Differential Equations, Springer, New York, NY, USA, 1993.

[22] M. Crampin and D.J. Saunders, ”On null Lagrangians”, Diff. Geom. Appl., 22, 131–146, 2005.

[23] D.J. Saunders, ”On null Lagrangians”, Math. Slovaca, 65, 1063–1078, 2015.

[24] A.J. Lotka, Elements of physical biology, Baltimore, 1925.

[25] V. Volterra, ”Flucations in the abundance of a species considered mathematically”, Nature, 18, 1–42, 1926.

[26] P.F. Verhulst, ”Notice sur la loi que la population poursuit dans son accroissement”, Correspondance mathématique et physique 10, 113–21, 1838.

[27] B. Gompertz, ”On the nature of the function expressive of the law of human mortality, and on a new method of determining the value of life contingencies”, Phil Trans Roy Soc. 27, 513–85, 1825.

[28] V.P. Collins, R.K. Loeffler, H. Tivey, ”Observations on growth rates of human tumors”, Am J Roentgenol Radium Ther Nuc Med. 78, 988–1000, 1956.

[29] W.O. Kermack and A.G. McKendrick, ”A contribution to the mathematical theory of Epidemics”, Proc. Roy. Soc. Lond. A, 115, 700–721, 1927.

